# Irreversible sterility of workers and high-volume egg production by queens in the stingless bee *Tetragonula carbonaria*

**DOI:** 10.1101/2020.06.05.136002

**Authors:** Francisco Garcia Bulle Bueno, Rosalyn Gloag, Tanya Latty, Isobel Ronai

## Abstract

Social insect reproduction is characterised by a division of labour. Typically, the queen is the sole reproductive female in the colony and the female workers are non-reproductive. However, in the majority of social insect species the workers are only facultatively sterile and remain capable of laying eggs under some conditions, such as when the queen dies. The Australian stingless bee *Tetragonula carbonaria* is noteworthy as workers never lay eggs, even if a colony loses its queen. Here we describe the reproductive anatomy of *T. carbonaria* workers (deactivated ovaries), virgin queens (semi-activated ovaries), and mated queens (activated ovaries). *T. carbonaria* mated queens have high-volume egg production compared to other female insects as each of their eight ovarioles (filaments of the ovary) produces approximately 40 eggs per day. We then conduct the first experimental test of absolute worker sterility in the social insects. Using a controlled microcolony environment, we investigate whether the reproductive capacity of adult workers can be rescued by manipulating the workers’ social environment (separating them from a queen) and diet (feeding them unrestricted highly nutritious honey bee royal jelly), both conditions which cause ovary activation in bee species where workers are facultatively sterile. The ovaries of *T. carbonaria* workers that are queenless and fed royal jelly remain non-functional, indicating they are irreversibly sterile and that ovary degeneration is fixed prior to adulthood. We suggest that *T. carbonaria* might have evolved absolute worker sterility because colonies under natural conditions are unlikely to ever be queenless.

## INTRODUCTION

In social insects the reproductive labour is divided between queens and workers (Hammond and Keller, 2004; Naug and Wenzel, 2006). The queen is typically the sole reproductive female and her specialised function in the colony is to lay eggs. Whereas, the non-reproductive female workers forage for food, defend the colony against predators and care for the queen’s brood (Michener, 1974).

In the majority of social insect species, workers are only facultatively sterile and have the ability to lay unfertilised haploid eggs that develop as males (Ebie et al., 2015; Khila and Abouheif, 2010; Wenseleers et al., 2004). The contribution of workers to the colony’s male output is highly variable across social insect species, and is determined by kinship-mediated cooperation and within-colony conflict over male production (Alves et al., 2009; Herbers, 1984; Mehdiabadi et al., 2003; Tóth et al., 2002; van Veen et al., 1990). Facultative worker sterility means that if the queen is lost and the colony cannot requeen, then workers can secure some reproductive success by producing males (Bourke, 1988).

The key determinants of the reproductive capacity of social insect workers is their nutritional status and social environment (Ronai et al., 2016). First, the presence of the queen, through aggressive interactions or pheromones, triggers programmed cell death in the ovaries of the workers (Luna-Lucena et al., 2018; Ronai et al., 2015); workers may therefore develop activated ovaries only in the queen’s absence. Second, ovary development is trophogenic (Lisboa et al., 2005) as the quantity or quality of nutrients has a strong effect on ovary development (Boleli et al., 2000; Boleli et al., 1999; Corona et al., 2016; Cruz-Landim, 2000; Jarau et al., 2010; Luna-Lucena et al., 2018; Schwander et al., 2010; Tambasco, 1975). For example, in the honey bee *Apis mellifera*, females fed highly nutritious royal jelly as adults have activated ovaries even in the presence of queen pheromone (Kamakura, 2011; Kamakura, 2014; Lin and Winston, 1998; Pirk et al., 2010).

Stingless bees (Meliponini) are a species-diverse pantropical clade of social insect, comprising Old World and New World subclades that diverged 60-70 MYA (Michener, 1974; Rasmussen and Cameron, 2009; Roubik, 1995).The reproductive habits of stingless bee workers have been best studied in the New World subclade, where workers produce males under queenright conditions (Beig et al., 1985; Boleli et al., 2000; Contel and Kerr, 1976), produce males only when queenless (Cruz-Landim, 2000; Tóth et al., 2004), lay trophic eggs which serve as nourishment for the queen (Wille, 1983), or never lay eggs (Boleli et al., 1999; Staurengo da Cunha, 1979). However, the reproductive habits of workers in the Old World subclade are relatively unknown.

In this study, we describe the reproductive anatomy of mated queens, virgin queens and female workers of the Australian stingless bee *Tetragonula carbonaria*. A colony of this Old World subclade species has one mated queen, thousands of non-reproductive female workers, some males and a few virgin queens (Gloag et al., 2007; Heard, 2016; Nunes et al., 2015). Notably, when the queen is removed from the colony, *T. carbonaria* female workers do not activate their ovaries (Nunes et al., 2015), suggesting they may be irreversibly sterile. To confirm this, we test whether manipulating the nutritional and social environment of age-matched adult female workers in a controlled microcolony environment can rescue their reproductive capacity (i.e. stimulate oogenesis). We hypothesised that if adult *T. carbonaria* workers are irreversibly sterile, then those fed an unrestricted highly nutritious diet in the absence of a queen would show no change in reproductive capacity.

## MATERIALS AND METHODS

### Biological material

Our study was carried out at the University of Sydney (Sydney, Australia) in December-January of 2017 and December-February of 2019. As the source colonies, we used seven *T. carbonaria* colonies (Colony 1–7) housed in wooden OATH hives (Heard, 2016). These colonies were queenright, weighed above average (8–9 kg, (Heard, 2016)) and had high forager activity at the entrance of the colony (∼100 bees exiting per minute).

To obtain age-matched workers we extracted two discs of late stage worker brood comb (approximately 200 bees per disk) from a source colony and placed the discs in a container. The containers were then placed in an incubator (34.5 °C) for approximately 5 days. *T. carbonaria* worker brood comb contains both male and female brood in identical cells. Any males that emerged were removed from the container.

### Worker and queen reproductive capacity

To assess the differences in reproductive anatomy between castes under natural conditions, we collected newly emerged workers (*n* = 200, per colony), recognised by their pale colour, from four of the source colonies (Colony 1–4), colour-marked their thorax using POSCA Pens and returned them to their source colony. After 14 days, the majority of marked workers have lost their mark or died, but we collected 10 marked workers from each source colony.

We collected the mated queen (unknown age) and one virgin queen (unknown age) from each of these four source colonies (Colony 1–4). To extract the queens, we set up hives with a plastic lid so that we could observe the brood comb. Once the mated queen was collected, we waited 24 h for a virgin queen to emerge on the brood comb and then extracted a virgin queen. These virgin queens were walking on top of the brood comb flapping their wings and were being fed via trophallaxis by the workers. All collected workers and queens were immediately placed into a −80 °C freezer.

### Manipulation of workers in microcolonies

To manipulate the social environment and diet of adult workers we designed laboratory-based microcolonies (see Fig. S1) using small plastic containers. Each microcolony was provided with a pellet of propolis (a mix of wax and plant resins that stingless bees use as nest building material (Heard, 2016)) and a pellet of pot-pollen (pollen is stored in pots made of propolis (Heard, 2016) and is the main protein source for stingless bees), both obtained from *T. carbonaria* colonies. Into each microcolony we placed newly emerged workers (*n* = 50 per microcolony, 8 microcolonies) from one of three source colonies (Colony 5–7). The eight microcolonies contained no queen and also no brood.

Four of the microcolonies were randomly assigned a control diet consisting of stingless bee honey and the other four paired (from the same source colony) microcolonies were assigned a royal jelly diet consisting of 50% stingless bee honey to 50% frozen royal jelly (Royal Jelly, Australia). Royal jelly is high in protein and contains important fatty acids(Altaye et al., 2010) and is fed to honey bee queens throughout their lifespan (Howe et al., 1985; Melampy and Jones, 1939; Viuda-Martos et al., 2008). Notably, feeding royal jelly to adult honey bee workers causes them to activate their ovaries (Cardoso-Júnior et al., 2020; Lin and Winston, 1998; Wang et al., 2014; Yang et al., 2017) and feeding it to female fruit flies, crickets and silkworms increases their fecundity (Hodin, 2009; Hodin and Riddiford, 2000; Kamakura, 2014; Kayashima et al., 2012; Kunugi and Mohammed Ali, 2019; Miyashita et al., 2016; Xin et al., 2016).

The microcolonies were checked daily. Food and water were replenished when needed. Any dead workers were removed and recorded (see Table S1).

From each microcolony we collected workers (*n* = 2) at 0 days of age. The remaining workers were collected at 14 days of age. Collected workers were immediately placed into a - 80 °C freezer.

### Ovary dissections

We extracted the paired ovaries from each bee collected from both natural colonies and microcolonies. The tiny ovaries of the workers need to be carefully dissected as they are fragile and likely to break. We placed the ovaries in a drop of distilled water on a microscope slide and covered them with a cover slip. The ovaries were imaged under a dissecting microscope (Leica IC80 HD) at 0.67X (mated queens and virgin queens) and 4X (workers) magnification. We analysed the images with ImageJ (Rasband, 1997) while blind to treatment.

We evaluated the reproductive capacity and anatomy of *T. carbonaria* females following the protocols used in our previous studies of honey bee ovaries (Ronai et al., 2017; Ronai et al., 2015). For each bee, we assessed the ovary activation state (deactivated, semi-activated or activated) and counted the number of ovarioles (filaments) in an ovary. We then quantified the length of the ovary as a proxy for the amount of ovary development or degeneration. For workers, we measured the length of both ovaries (from the tip of the longest ovariole of each ovary to where the two oviducts join, Fig. S2A). For mated queens and virgin queens, it was impossible to measure the length of the ovarioles as they are tightly curled, so we measured the elliptical area of the ovary (area = *a x b x* π, where *a* is the major diameter divided by 2 and *b* is the minor diameter divided by 2, Fig. S2B). If still attached after dissection, we noted the presence and status (semen present or absent) of the spherical spermatheca, the specialised organ used to store the sperm, and measured its diameter if present (Fig. S2).

For all workers, we investigated whether the length of the right and left ovary was correlated using a Pearson correlation so we could then compare the length of the longest ovary in our experimental treatments. To compare the length of the longest ovary of workers fed a Control diet and Royal jelly diet, we conducted a Student t-test (all assumptions were met) using R Studio (Version 1.3.959), https://www.rstudio.com/), with R packages the function “t.test” in the R package *stats v4.0.0* (Team, 2013).

## RESULTS

### Reproductive morphology of workers in queenright colonies

Queenright 14-day-old *T. carbonaria* workers had deactivated ovaries as no developing oocytes were present inside the ovarioles (Fig. 1). The number of ovarioles in the ovary of workers varied from four ovarioles to no ovarioles. The mean length of the longest ovary was 0.65 mm ± 0.15 mm (*n* = 40, Table S2). The length of the right and left ovary was asymmetric (*P* =0.098, Table S3). Only 20% of workers (*n* = 8) had an identifiable, but vestigial spermatheca with a mean diameter of 0.05 mm ± 0.01 mm (Fig. 2A, Table S2).

**Figure 1.**
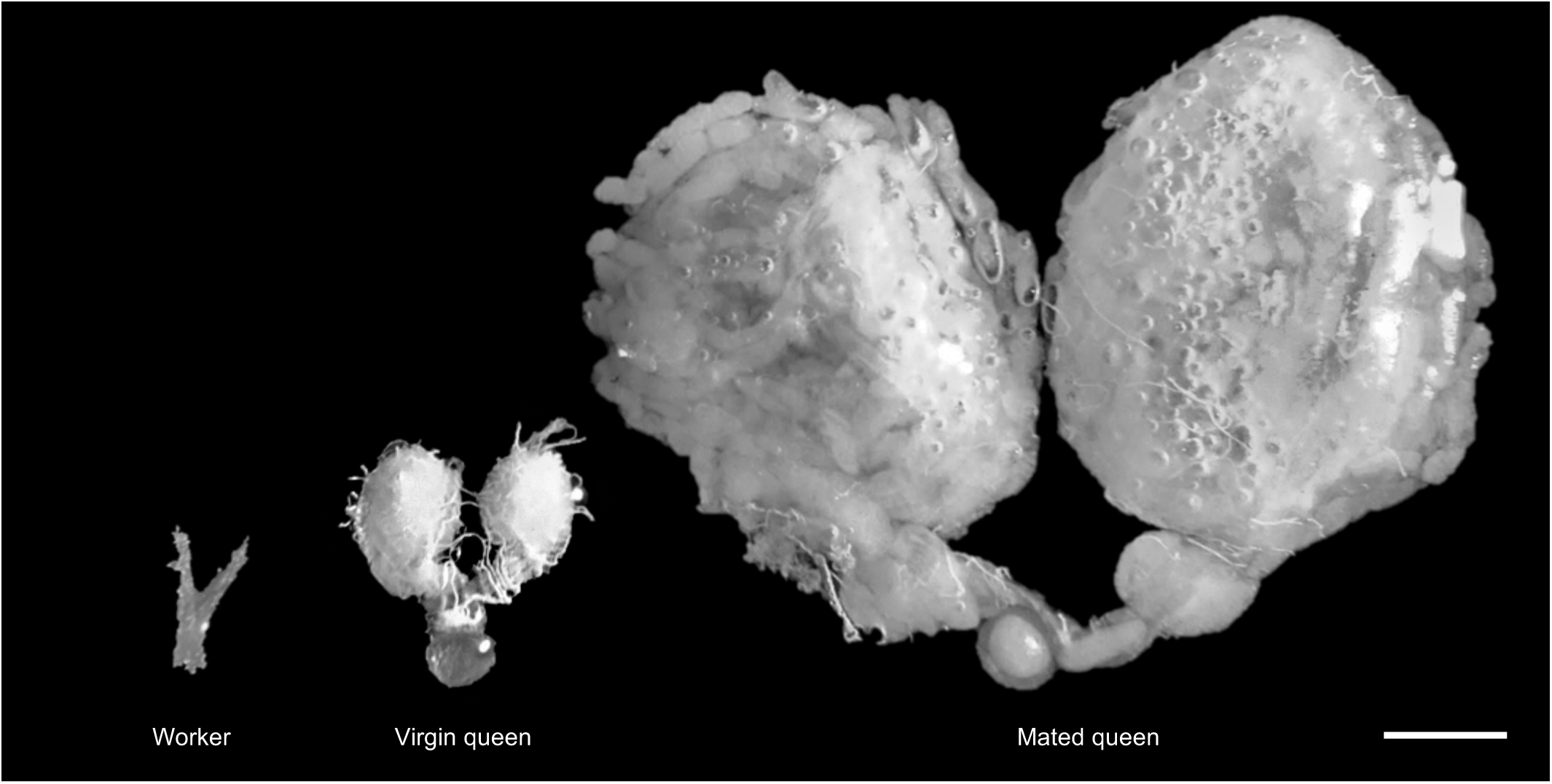
The ovaries of *T. carbonaria* females. Queenright 14-day old workers have deactivated ovaries (no developing oocytes present) with zero to four ovarioles. Virgin queens have semi-activated ovaries (oocytes not mature) with four ovarioles, Mated queens have activated ovaries (mature oocytes present) with four ovarioles. Scale bar 1 mm.

**Figure 2.**
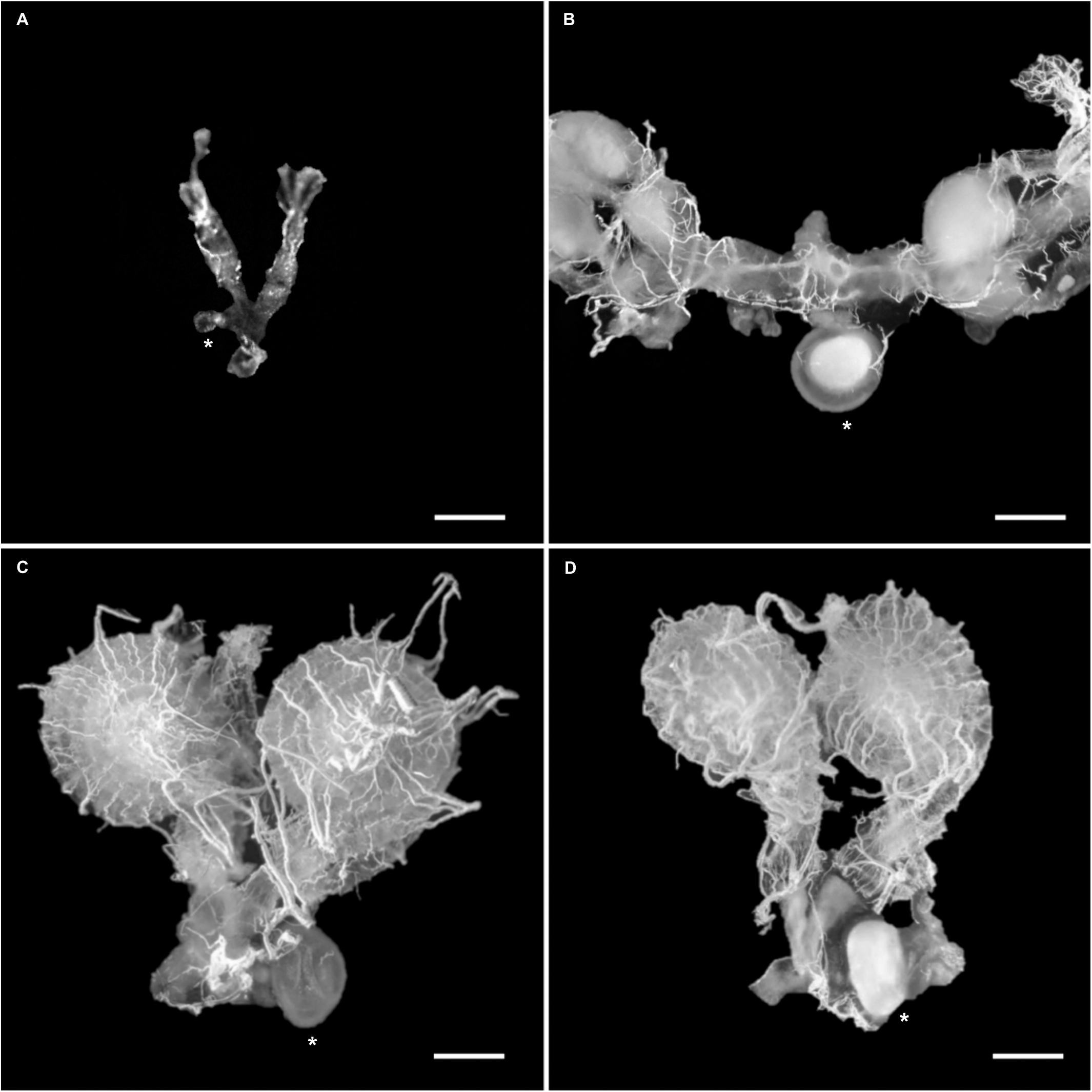
The spermatheca of *T. carbonaria* females. A) Queenright 14-day old worker with a vestigial spermatheca. B) Mated queen with a full spermatheca reservoir containing semen. C) Virgin queen with an empty spermatheca reservoir. D) Virgin queen with a full spermatheca reservoir containing semen and semi-activated ovaries, suggesting mating occurred recently. Spermatheca (*). Scale bars 0.5 mm.

### Reproductive morphology of virgin queens and mated queens

*T. carbonaria* queens had four ovarioles per ovary (Fig. 2). We observed that the ovaries of *T. carbonaria* are meroistic polytrophic (Büning, 1994; Martins and Serrão, 2004) as a single germ cell produces both an oocyte and connected nurse cells, which move with the developing oocyte through the ovariole. As an oocyte moves from the tip of the ovariole to the base it progresses through stages of development (stem cell, germarium and vitellarium) (Ronai et al., 2015).

Virgin queens had semi-activated ovaries with oogenesis underway (Fig. 1) and the mean area of the largest ovary was 1.02 mm^2^ ± 0.12 mm^2^ (*n* = 4, Table S4). The spermatheca had a mean diameter of 0.55 mm ± 0.03 mm (n = 3, Table S4). For two of the virgin queens the spermatheca reservoir was transparent indicating it was empty (Fig. 2C), however the third virgin queen had a filled spermatheca reservoir indicating that she mated within the 24 hours that there was no mated queen in the colony (Fig. 2D).

Mated queens had activated ovaries full of mature oocytes (Fig. 1). Some of the activated ovaries had yellow bodies present, a consequence of previous oviposition (Feneron and Billen, 1996; Gobin et al., 1998). The mean area of the largest ovary was 13.41 mm^2^ ± 2.99 mm^2^ (*n* = 4, Table S4), approximately thirteen times larger than the virgin queens. In order to fit inside the abdomen (approximately 6.5 mm in length) of the queen the ovaries are curled tightly in a spiral and occupy the majority of the abdominal cavity. The spermatheca had a mean diameter of 1.05 mm ± 0.81 mm (*n* = 2, Table S4) and for both the reservoir was filled with a milky white substance indicating the presence of semen (Fig. 2B).

### Reproductive morphology of queenless workers fed royal jelly diet and control diet

Both royal jelly diet and control diet queenless workers had degenerated and deactivated ovaries. The mean length of the longest ovary of 0-day old workers was 0.56 mm ± 0.1 mm (*n* = 16, Table S2). The mean length of the longest ovary of 14-day-old workers fed a control diet was 0.57 mm ± 0.14 mm (*n* = 40, Table S2) and a royal diet was 0.59 mm ± 0.11 mm (*n* = 40, Table S2). The mean length of the right and left ovary was significantly correlated for both control diet and royal diet workers (*P* = 0.001 and 0.007 respectively, Table S3). There was no significant difference in the mean length of the longest ovary between the control diet and royal diet treatments after 14 days (*t* = −0.68194, *df* = 75.203, *p* = 0.4974, Fig. 3).

**Figure 3.**
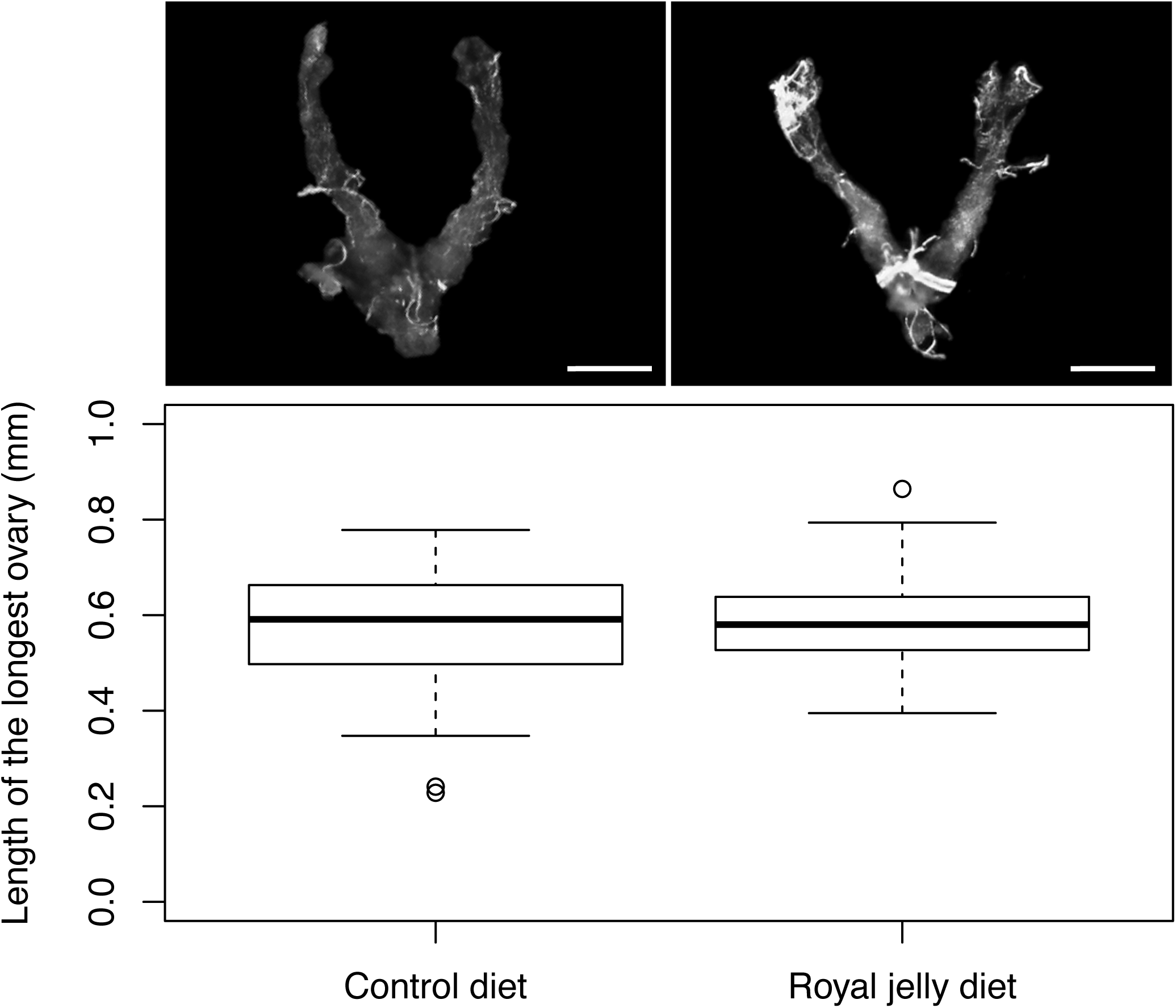
Ovary length of 14-day old queenless *T. carbonaria* workers. The workers were fed a Control diet (*n* = 40, 10 per microcolony) or a Royal jelly diet (*n* = 40, 10 per microcolony). Scale bars 0.25 mm. There was no significant difference between the treatment groups.

## DISCUSSION

An unrestricted high quality diet and the absence of the queen failed to rescue the reproductive capacity of *T. carbonaria* adult workers, indicating that they are irreversibly sterile. This absolute sterility is consistent with observations from natural nests in which workers never lay eggs, even if queenless (Gloag et al., 2007; Nunes et al., 2014). Absolute sterility of workers is relatively rare among social insect species. Only 13 genera have so far been classified as having truly sterile workers based on observations under natural conditions (Ronai et al., 2016); whether sterility is irreversible in these species has not yet been tested experimentally.

Why have a few genera of social insect evolved absolute worker sterility and forgone the obvious benefits of facultative sterility? It is likely that the life history of some species predisposes them to the evolution of absolute worker sterility. *T. carbonaria* have a high and constant rate of queen production (Heard, 2016) when compared to other stingless bee species with facultative worker sterility (Grosso et al., 2000; Ribeiro et al., 2003; Van Veen and Sommeijer, 2000). A colony is unlikely to ever have no queen present and there would never be a need for workers to lay eggs (Engels and Imperatriz-Fonseca, 1990; Michener, 1974).

Species that have a constant supply of virgin queens from which they can select the most fecund one or rear new queens very quickly may be more likely to evolve absolute worker sterility (Sommeijer et al., 1994). Another possible explanation for the evolution of absolute worker sterility is the high rate of colony-takeovers in *T. carbonaria* (Cunningham et al., 2014; Gloag et al., 2008). This behaviour reduces the chance that a colony would ever have a prolonged period where it was queenless and that the workers would need to lay eggs.

Alternatively, absolute worker sterility may resolve reproductive conflict within colonies. *Tetragonula carbonaria* is a monogamous species (Green and Oldroyd, 2002). Under monogamous queens, workers in a colony are all full sisters and workers would be more related to their own sons (*r = 0.5*) than the sons of the queen (*r = 0.25*). If workers have the ability to lay eggs, there would be worker-queen conflict for male production in the colony (Trivers and Hare, 1976). The evolution of absolute worker sterility in monogamous species eliminates the worker-queen conflict to the benefit of the queen. However, many social insects, including most New World sublclade stingless bees, have both monogamous queens and workers that lay eggs (Vollet-Neto et al., 2018).

We find that *T. carbonaria* workers emerge from pupation with non-functional ovaries that contain no germ cells. The degeneration of the ovary in *T. carbonaria* workers must therefore occur prior to adulthood. To date, only one other genera of stingless bee in the New World subclade has been proposed to have absolute worker sterility, *Frieseomelitta* spp. (Boleli et al., 2000; Boleli et al., 1999; Luna-Lucena et al., 2018; Ronai et al., 2016) and their ovaries degenerate during the last pupal stage via apoptosis (Boleli et al., 1999). Further work is needed to establish whether the degeneration of the ovaries in *T. carbonaria* workers is initiated during the embryonic, larval or pupal stage. In *T. carbonaria* the ovary degeneration in workers is likely determined by their nutritional environment pre-emergence as worker-destined larvae provisioned with a smaller quantity of food than queen-destined larvae (Nunes et al., 2015).

Even when the workers of social insects have activated ovaries, they may not have a storage organ for semen. The spermatheca is thus normally absent from workers (Gobin et al., 2008; Hölldobler and Wilson, 1990; Khila and Abouheif, 2010; Van Eeckhoven and Duncan, 2020). However, *T. carbonaria* workers sometimes have a vestigial spermatheca, the first reported case in the Old World subclade of stingless bees. Vestigial spermatheca are present in workers of the one other stingless bee genera with absolute worker sterility (Boleli et al., 2000) and also in honey bee species with facultative worker sterility (Gobin et al., 2006; Gotoh et al., 2013). The presence of a vestigial spermatheca in species where the workers have degenerated ovaries provides evidence that spermatheca degeneration is a separate process to the degeneration of the ovaries.

Our analyses of ovary anatomy in *T. carbonaria* highlights that the exceptionally high reproductive capacity of queens is achieved in different ways in the different clades of eusocial bees. Stingless bees and honey bees (*Apis* spp.) each evolved highly eusocial life histories independently, from an ancestral corbiculae apid with more simple sociality (Cardinal and Danforth, 2011). *Tetragonula carbonaria* queens lay approximately 300 eggs per day (Heard, 2016) but have only 8 ovarioles in total; therefore each ovariole is elongated and produces around 37 eggs per day. In contrast, *A. mellifera* queens lay approximately 2000 eggs per day (Page and Erickson, 1988) but have on average 320 ovarioles in total (Jackson et al., 2011); therefore each ovariole is relatively short and produces only around 6 eggs per day. That is, the lower number of ovarioles in stingless bee queens when compared to honey bee queens is compensated by high volume egg production in each ovariole of the ovary.

In conclusion, *T. carbonaria* workers are irreversibly sterile as adults. Conducting similar studies of workers in the other *Tetragonula* spp. would establish whether absolute sterility is a characteristic of this genus. Studies of absolute worker sterility, and the factors that favour its evolution, are likely to provide a deeper understanding of the evolution of eusociality.

## ACKNOWLEDGEMENTS

We thank Liz Gibson and Peter Clarke for their support in lending colonies of *T. carbonaria*.

## AUTHOR CONTRIBUTIONS

Conceptualisation: F.G.B.B, I.R.; Methodology: F.G.B.B, I.R.; Validation: F.G.B.B, I.R.; Formal analysis: F.G.B.B, I.R.; Investigation: F.G.B.B; Writing - original draft: F.G.B.B, I.R.; Writing - review & editing: F.G.B.B, R.G., T. L., I.R.; Visualisation: F.G.B.B, I.R.; Funding acquisition: R.G., T.L..

